# Synthetic Photoplethysmography (PPG) of the radial artery through parallelized Monte Carlo and its correlation to Body Mass Index (BMI)

**DOI:** 10.1101/2020.07.22.214015

**Authors:** Tananant Boonya-ananta, Andres J. Rodriguez, Ajmal Ajmal, Vinh Nguyen Du Le, Anders K. Hansen, Joshua D. Hutcheson, Jessica C. Ramella-Roman

## Abstract

Cardiovascular disease is one of the leading causes of death in the United States and obesity significantly increases the risk of cardiovascular disease. The measurement of blood pressure (BP) is critical in monitoring and managing cardiovascular disease hence new wearable devices are being developed to make the BP metric mode accessible to physicians and patients. Several wearables utilize photoplethysmography from the wrist vasculature to derive BP assessment although many of these devices are still at the experimental stage. With the ultimate goal of supporting instrument development, we have developed a model the photoplethysmographic waveform derived from the radial artery at the volar surface of the wrist. To do so we have utilized the relation between vessel biomechanics through Finite Element Method and Monte Carlo light transport model. The model shows similar features to that seen in PPG waveform captured using an off the shelf device. We observe the influence of body mass index (BMI) on the PPG signal. A degradation the PPG signal of up to 40% in AC to DC signal ratio was thus observed.

## 1. Introduction

Elevated blood pressure (BP) is considered one of the highest risk factors for cardiovascular disease. In fact, it has been estimated that 47% of all coronary heart disease worldwide is attributable to high BP^1^. BP is less than 120/80 mmHg in normotensive individuals, between 120-129/80 mmHg in prehypertensive individuals, between 130-139/80-89 mmHg in hypertensive stage 1, above 140/90 mmHg in hypertensive stage 2, and above 180/120 mmHg in hypertensive^2^. In adults, the risk of cardiovascular disease is significantly increased with obesity^3-5^, defined as having a BMI of over 30 kg/m^2^ ^6-9^. BMI is found to correlate strongly with increased 24 hour blood pressure as well as non-dipping nocturnal blood pressure^10,11^. Physiological changes that occur in individuals with obesity may increase the measurement uncertainties, and optical means for monitoring blood pressure will have to account for obesity-related changes to the skin optical properties ^12^ and increased subcutaneous tissue thickness.

There are several ways to evaluate BP. Sphygmomanometers and oscillometric devices are standard techniques^13^, and the most common clinical method of measuring blood pressure is through auscultations using a sphygmomanometer with the arm maintained at heart level^14^. Invasive arterial line blood pressure measurement is often used in the Intensive Care Unit (ICU), where an arterial catheter is placed inside the radial artery of a patient to directly measure vessel internal pressure^15,16^.

Research into cuff-less and continuous BP devices based on photoplethysmography (PPG) is rapidly expanding. These systems provide unique diagnostic opportunities, such as monitoring of nocturnal hypertension^10^ a condition strongly associated with cardiovascular events and organ damage. The PPG waveform can provide information such as pulse wave velocity, the relative change in blood volume and pressure wave reflection^17^. The systolic peak and diastolic peak are the primary and secondary features respectively as seen in Figure 1. The primary feature is directly representative of systolic performance during the cardiac cycle^17,18^. The vessel dilates due to the pressure wave generated by ventricular contraction. The pressure profile inside the arteries changes as the pressure wave propagates downstream from the heart to peripheral vasculature^19-21^. This is due to vessel compliance, vessel branch and bifurcations, and wave reflections encountered by the pulse during its journey from the heart. The dicrotic notch moves further away from the systolic peak as arterial line measurement location moves further down the arterial tree.

**Figure 1.**
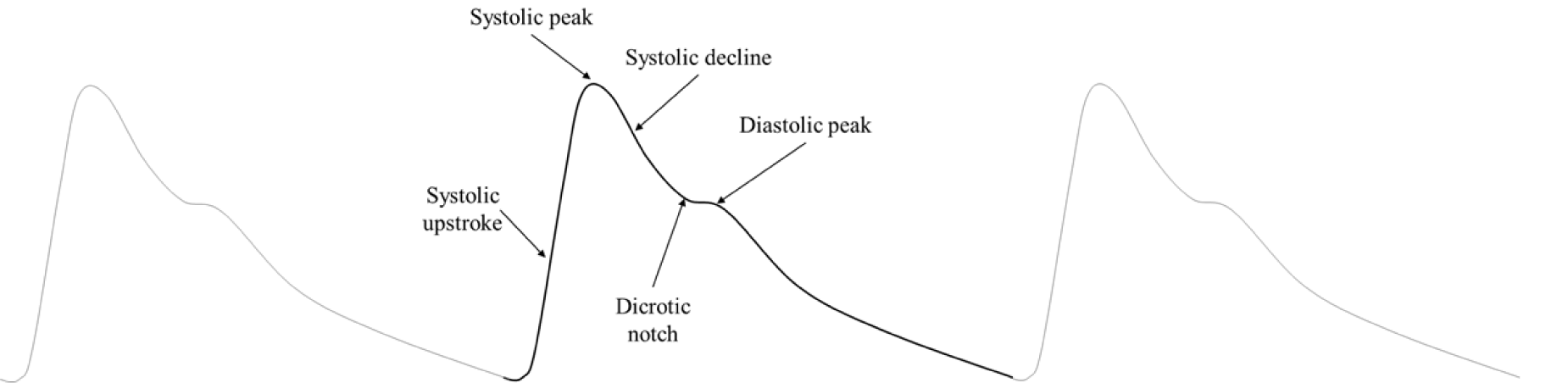
Typical PPG waveform with constitutive components. PPG wave components shows the systolic upstroke to systolic peak, then systolic decline into the dicrotic notch and the diastolic peak.

Measurement of blood pressure at the radial artery located at the wrist is desirable since it is easily accessible, and instrumentation can be developed for a wearable device. The blood pressure in the radial artery at the volar surface of the wrist is more representative of blood pressure in the main arterial network as opposed to pressure in superficial vasculature of the skin^22^.

In order to derive a better understanding of devices based on PPG, several groups have conducted light propagation models using Monte Carlo (MC) simulations^23-25^. Yet to our knowledge, these studies have not attempted to capture the PPG waveform in its entirety and have focused on an idealized skin model.

The design of wearable devices must account for the population diversity, including physiological changes due to obesity^26-28^ as well as skin tones.

Our study has two main goals. First, we develop a more realistic representation of light interaction with the radial artery during a pulse as seen by a commercially available photoplethysmographer (Nellcor™ by Covidien). This location and instrumentation are the focus of our experimental efforts toward developing low-cost cuffless BP sensors. Second, we use our model to study the influence of increased BMI on the PPG waveform.

Recent studies have shown that obesity creates variations in skin physiology^5,29,30^. These changes include but are not limited to skin barrier function, epidermal changes, dermal changes, and changes to the vascular and capillary recruitment function. In our model, both skin tone and BMI diversity will be considered.

## 2. Material and Methods

The behavior of arterial pulsatile flow can be modeled as a pulsatile flow through an elastic walled tubing^19^. The key distinction between rigid versus elastic walled flow is the fluid streamline velocity profile (Figure 2A, 2B). The typical Poiseuille velocity profile for ideal, fully developed fluid flow in a rigid tube is constant along the length of the tube (Figure 2A), whereas in elastic-walled tubing, the velocity profile is dependent upon the location along the streamline (Figure 2B). In elastic tubes, pressure waves cause a local change in fluid pressure that is then propagated downstream. Wall compliance results in oscillation of the wall and changes to tube diameter with induced pulsatile flow, as seen in the vascular system.

**Figure 2.**
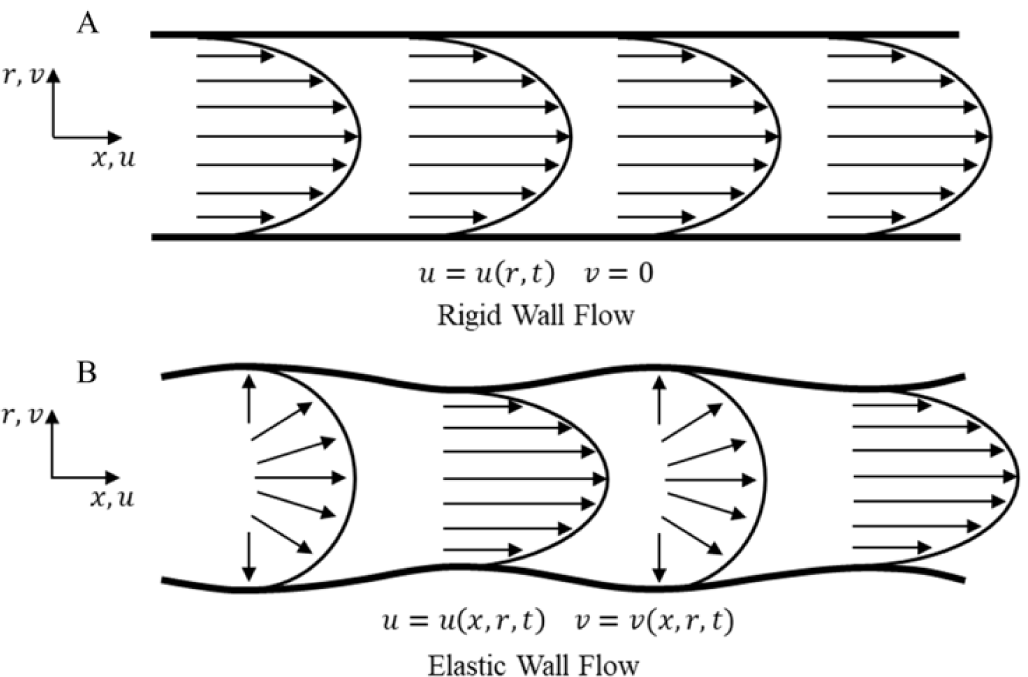
Fluid flow in rigid and elastic tubing (adapted^19^). (A) The top diagram shows fully developed velocity profile for flow in a rigid walled tube. (B) Second diagram shows flow in an elastic walled tube.

The mechanics of large vessels such as the aorta, carotid artery, and large coronary arteries has been studied extensively^31-36^ but several authors have noted that smaller peripheral vessels, such as the one under study here, exhibit similar mechanical behaviors to larger ones^37,38^. Considering the biomechanical changes in the arterial vascular bed, our model of PPG is divided in two parts. First, a finite element model is used to observe the dilation of the radial artery under applied pressure. Then, a Monte Carlo model is applied to this dynamic geometry to observe loss of optical signal in relation to the pulse.

### Vessel Geometry

We have used Finite Element Analysis (FEA) to model the mechanical behavior of an arterial vessel. Our model allows us to visualize the motion of an arterial wall under an applied internal pressure. The artery is modeled in Solidworks computer-aided design (CAD) software. We have chosen the volar surface of the distal forearm as the location of our virtual PPG instrument. This choice is governed by our interest in the development of wrist-based wearable devices targeting that vessel.

A drawing of the vessel with associated dimensions is shown in Figure 3A. The radial artery has been reported to have 2.5mm inner diameter and 0.2 wall thickness^38-40^. Vessel wall mass density and Poisson’s ratio were set to 1160 kg/m^3^ and 0.49^36,38,41^ respectively. A radial artery wall Young’s modulus of 0.70MPa was used^38^. Arterial mechanical stiffness ranges between 0.70MPa to 1.10MPa^38^.

**Figure 3.**
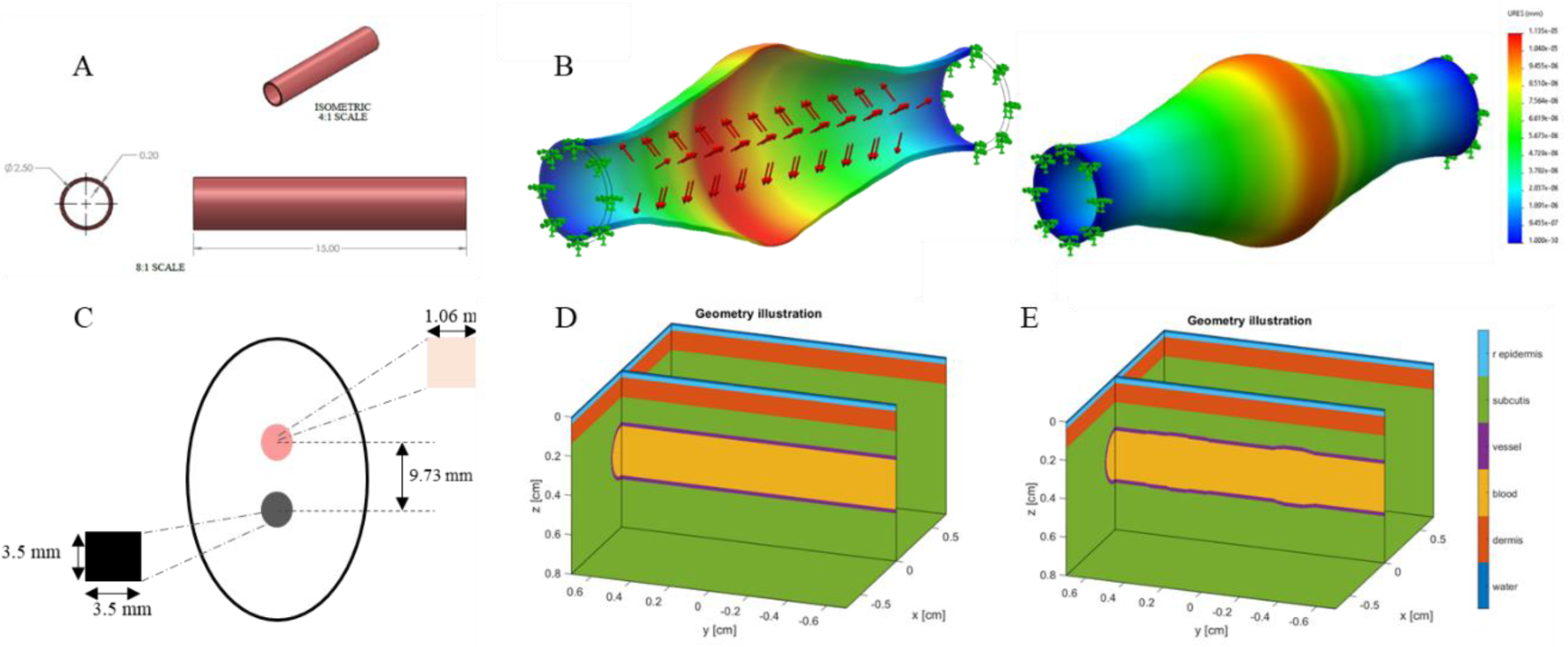
Vessel 2D CAD geometry drawing. This is the model used for the radial artery to understand its behavior during the cardiac cycle. All dimensions in [mm]. (B) Arterial wall dilation contour plot of total displacement. Red regions show regions of highest displacement and blue for lowest displacement. Note depicted displacement scale is exaggerated. (C) Patch sensor schematic for adaptation to Monte Carlo. (D) Base geometric configuration for Monte Carlo Model. The various layers represented by a specific color developed in the Monte Carlo geometry. (E) Illustration of pulse shape generating features in Monte Carlo. Geometric parameters of arterial dilation and pressure wave propagation is adapted into Monte Carlo through variation in geometry of the artery. Pulse features are developed through Equations 1-3.

Radial artery pressure has been shown to be 5 to 15 mmHg^42^ higher than the brachial artery, where blood pressure is often measured using the sphygmomanometer. An internal pressure of 130 mmHg^42^ was imposed to the vessel representing the radial artery. The result of the FEM model at peak applied force is shown in Figure 3B. The figure shows a time snapshot of the vessel shape as a time-dependent pressure pulse propagates from the heart to the radial artery. This is different from a quasi-static pressure vessel representation often used to analyze vascular wall mechanics.

The arterial wall is often treated as incompressible^36,43^, pressure changes applied internally translates to vessel wall dilation, but the total wall volume is conserved. This incompressibility estimation is seen in the FEM model. Changes to the inner wall and outer wall radii are relatively small in comparison to total vessel dilation. Vessel walls have been reported to experience 10-20% diametral strain^44,45^. Applying 0.70MPa stiffness, vessel wall shows total dilation of 0.4 mm in diameter. The mechanical behavior of the artery during pulse propagation is integrated into the light transport model (Monte Carlo) through changes in the vessel geometry.

This was done by deriving the mathematical envelope of the FEA derived pulse geometry using a double ellipsoid shown in Equations 1-3.

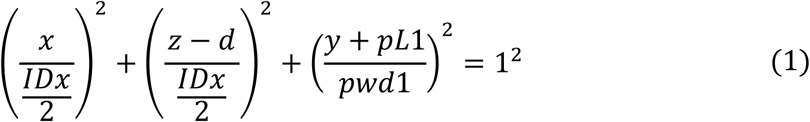

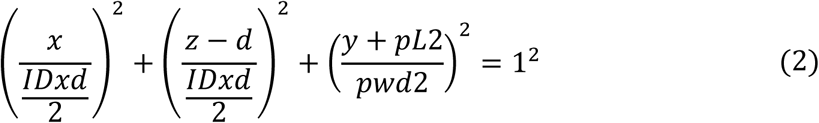

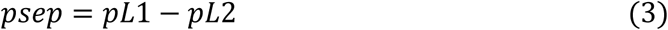

In Equations 1 and 2, *x, y*, and *z* represent each direction in the frame, *d* locates the ellipsoid in the vertical, *z*, direction in the media. In Equation 1, *IDx* defines the dilation of the inner diameter of the vessel, *pL*1 is the location of the primary ellipsoid in the *y*-axis, and *pwd*1 defines the width of the primary ellipsoid in the *y* direction. In Equation 2, *IDxd* represents the dilation of the inner diameter of the secondary ellipsoid, *pL*2 is the location of the secondary ellipsoid in the *y-axis*, and *pwd*2 defines the width of the secondary ellipsoid in the *y* direction. Equation 3 defines the *psep* parameter as the separation between the first and second ellipsoids. The geometry developed with these equations is shown in Figure 3E.

### Monte Carlo Model

Aside from the radial artery, the Monte Carlo simulation also considered three skin layers: epidermis, dermis and subcutaneous tissue with the radial artery inserted into the adipose tissue layer. The Monte Carlo model used in this work is an adaptation of MCMatlab developed by Marti, D et al^23^. MCMatlab converts S. Jacques’ “mcxyz.c”^24^ from C-based code to a compact tool usable through Matlab interface. Monte Carlo framework simulates a 3-layer model to include epidermis, dermis and subcutaneous tissue with the radial artery inserted into the adipose tissue layer. The epidermis consists of multiple sublayers with various components, including melanin and melanocytes^46^. The dermis is comprised of connective tissue along with blood vessels (capillaries and arterioles) with blood volume ratios ranging from 0.2% to 7% and water volume concentration around 65%^47,48^. The subcutis consists of subcutaneous adipose tissue and other connective tissue^49^.

In this work, we modeled a commercial PPG (Nellcor™ Covidien, this device also provide pulse oximetry). This is a clinical system that incorporates two wavelength sources at 660 nm peak 22 nm FWHM and 890nm peak 54nm FWHM. The sensor provides and effective beam waist radius of 0.06 cm. The detector is located 0.97 cm away from the sources providing reflectance PPG signal. Light collector area is measured to be 0.35cm x 0.35cm which is an effective collector area of 0.1225cm^2^ with numerical aperture of 0.866. The sensor schematic is shown in Figure 3C.

The geometry of our Monte Carlo simulation is shown in Figure 3D. The first layer mimics the epidermis. The thickness of human epidermis is approximately 0.10 mm^50,51^. To avoid using a very small voxel size in the simulation, we chose to scale the optical properties of the epidermis to match its optical thickness^52^.

The dermis layer was 1.0 mm in thickness. Below the dermis, a subcutaneous adipose tissue layer was added. Within this layer, the radial artery is modeled as a cylinder situated at 2.5mm depth from the top surface^39,40^. The target radial artery is constructed with a vessel wall and the internal lumen is filled with blood. Optical properties used in Monte Carlo simulations are extracted from previous work^47,53 25,54-57^.

Monte Carlo geometry frames a 1.4 × 1.4× 0.8 cm^3^ volume with 200 × 100 × 100 elements in each direction. Monte Carlo simulations were performed on Windows 10 64-bit Operating System with Intel® Core™ i7-8700 CPU 3.20GHz, NVIDIA GeForce GTX 1070 GPU, and 32GB RAM, and Windows 10 64-bit Operating System with Intel® Core™ i7-8750H CPU 2.20GHz, NVIDIA GeForce GTX 1050 Ti GPU, and 16GB RAM. Simulations speeds varied between two million photons per minute to 70 million photons per minute. Waveform simulations are conducted at 89 incremented pulse positions at 100 million to one billion photons each.

Analyses of the performance of commercial systems and functionality of arterial PPG must account for factors of skin tone and physiological changes with the development of obesity^26-28^. The melanin content of the skin increases with darkening skin tone. Melanin volume fraction variation can be used to model the various pigmentation levels. Lightly pigmented skin adults have a melanin volume fraction of 1.3-6.3%, moderately pigmented skin adults have a 11-16% melanin volume fraction, and darkly pigmented skin adults have an 18-43% melanin volume fraction in the epidermis^47^.

Dermal thickness variations with BMI is shown to range from 1.0mm to over 2.5mm thickness^50^. Along with changes to the skin, the artery itself is situated deeper in the subcutaneous adipose tissue as adipose tissue layer thickness with increasing level of obesity. The signal from the radial artery is discovered to vary in depth with different levels of obesity from 2.5mm for non-obese individuals at BMI of 25 to 3.5mm for BMI of 45^39,40^. Trans-epidermal water loss (TEWL) the process where water evaporates through the epidermis from the dermis is disruption in the obese^30^. As BMI increases so does TWEL^58^, and this effect directly contributed to dry skin and skin irritation often seen in the obese population. A summary of all the parameters utilized in the modeling is shown in Table 3.

**Table 1.** Arterial Wall Material Properties

| Material Property | Artery Wall |
| --- | --- |
| Young's Modulus (kPa) | 700 |
| Poisson's Ratio | 0.49 |
| Mass Density (kg/m<sup>3</sup>) | 1160 |
| Mesh Elements | 52610 |

**Table 2.** Optical properties at two corresponding wavelengths

| Layer | $\mu_a [cm^{-1}]$ | $\mu_s [cm^{-1}]$ | g | n |
| --- | --- | --- | --- | --- |
|  | 660nm/890nm | 660nm/890nm |  |  |
| rEpidermis | 0.3442/0.3184 | 121.2/224.7 | 0.9 | 1.4 |
| Dermis | 0.5453/2459 | 208.6/116.7 | 0.9 | 1.4 |
| Fat | 0.0001/0.0217 | 249.7/189.8 | 0.9 | 1.4 |
| Vessel Wall | 0.8/0.8 | 230/230 | 0.9 | 1.4 |
| Blood | 2.026/6.32 | 75.76/56.18 | 0.9 | 1.4 |

**Table 3.**
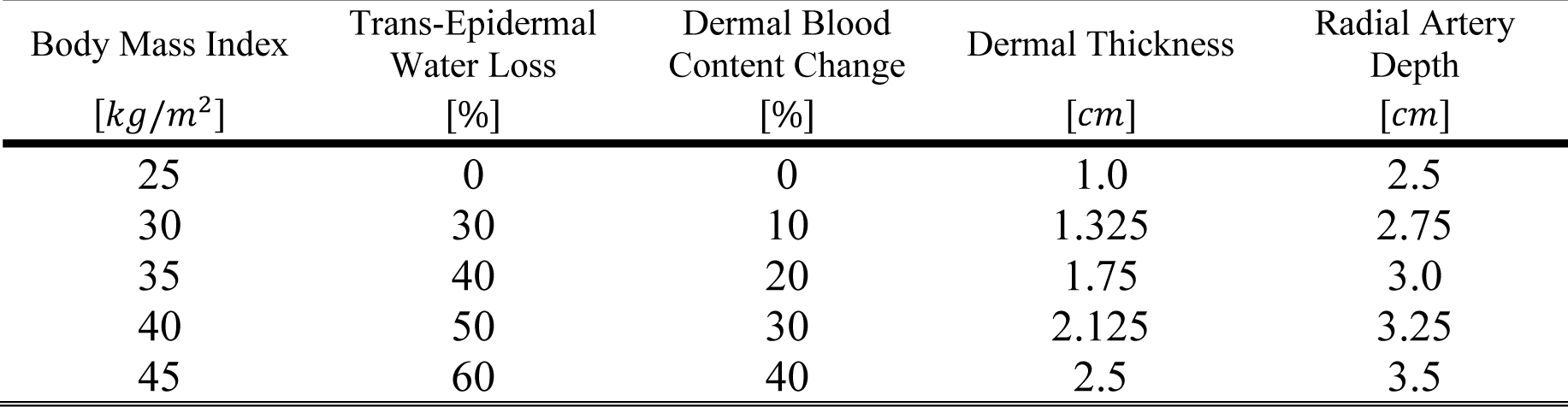
Skin changes simulation parameters attributed with increasing Body Mass Index and Obesity

| Body Mass Index<br>[$kg/m^2$] | Trans-Epidermal<br>Water Loss<br>[%] | Dermal Blood<br>Content Change<br>[%] | Dermal Thickness<br>[$cm$] | Radial Artery<br>Depth<br>[$cm$] |
| --- | --- | --- | --- | --- |
| 25 | 0 | 0 | 1.0 | 2.5 |
| 30 | 30 | 10 | 1.325 | 2.75 |
| 35 | 40 | 20 | 1.75 | 3.0 |
| 40 | 50 | 30 | 2.125 | 3.25 |
| 45 | 60 | 40 | 2.5 | 3.5 |

**Table 4.** Waveform baseline signal changes at different levels of obesity

| Waveform | Photons<br>[millions] | Baseline Change<br>[%] |
| --- | --- | --- |
| Obese 1 | 100 | 13.0 |
|  | 1000 | 12.7 |
| Obese 2 | 100 | 42.4 |
|  | 1000 | 36.5 |
| Obese 3 | 100 | 54.5 |
|  | 1000 | 52.4 |

Changes in Table 3 are applied to the MC geometry through two different methods. The changes to TWEL and dermal blood contented are applied to the media layer properties. TWEL changes the amount of water in the dermal layer. The water loss percentage is multiplied to standard dermal water content. Similarly, dermal blood content percentage changes are multiplied to blood content in the dermis. For geometric changes of dermal thickness and artery depth, these properties are changed directly in the geometry construction.

## 3. Results

A power absorption map of our geometry is shown in Figure 4A and Figure 4B. This is a single slice in the three-dimensional geometry at a single location of the pulse waves.

**Figure 4.**
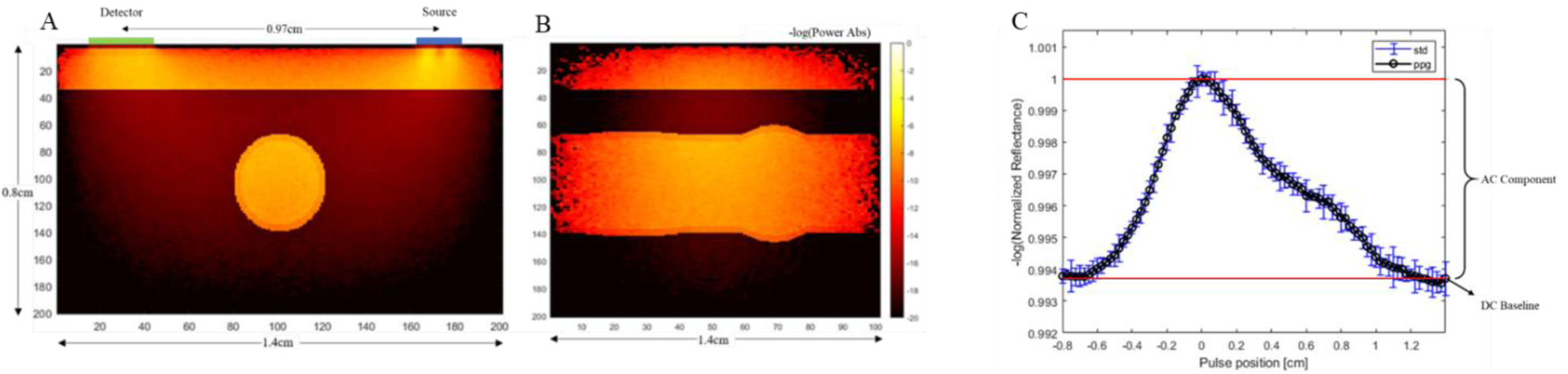
Monte Carlo slices of the negative logarithm of power absorbed color map of radial artery configuration at single pulse location at 660nm wavelength. A and B shows the front view and side view, respectively. (C) PPG Curve generated for non-obese case. PPG signal is developed through normalized reflectance signal collected at each pulse position.

Figure 4C shows an example of PPG waveform generated for the non-obese case. The returned signal is comprised of an AC and a DC baseline signal, where the AC signal represents the pulsatility of the vasculature. A comparison of the AC to DC signal ratio of each waveform shows the degradation of the signal with increasing BMI. Different PPG signals are generated through varying four different physiological changes observed with obesity. These changes can be seen in Table 2. Preliminary experimental data using the PPG sensor (Figure 5) shows features similar to that seen in simulation curve. The PPG of three different individuals was captured experimentally (Figure 5A). There is significant variation between individuals (Figure 5A green, red, blue), however the features of the shape can be identified in all three cases. The PPG data is adjusted from pulse shape in time to PPG with respect to pulse position (Figure 5B, 5C). This is done by matching the start and end point of the two sets as well as the primary peak and overlay on top of each other.

**Figure 5.**
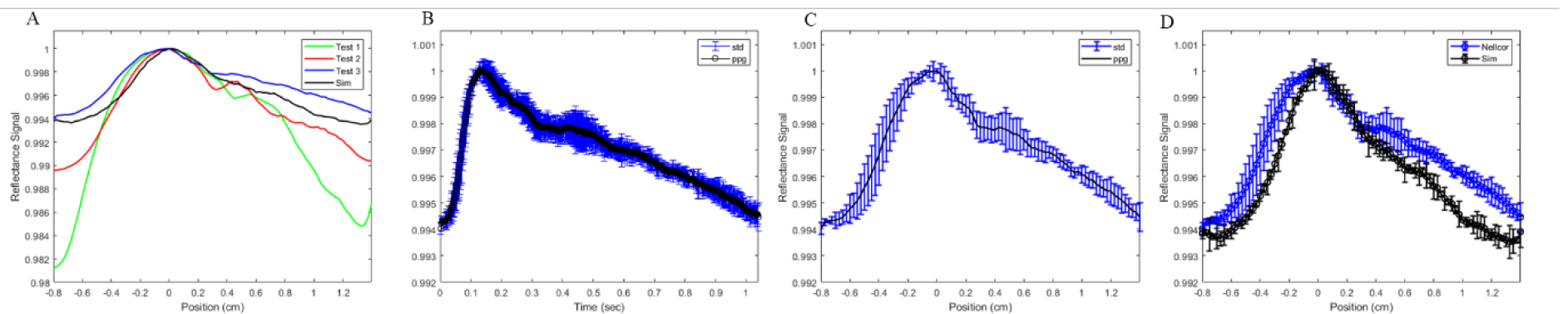
PPG data of radial artery taken at the volar location of the wrist of three different preliminary tests, each test is from a different individual. Note x-axis label is in time whereas simulation data pulse is developed through pulse position. (B) Experimental data of Test 3. (C) PPG adjusted for from time to pulse position. (D) Overlay of experimental PPG and simulated waveform.

Figure 6 shows changes AC/DC signal from four different changes attributed to obesity across different BMI levels. Figure 6A shows a 14.0% total signal decline associated with changes to trans-epidermal water loss between normal to highest BMI level. Figure 6B shows an 18.0% change with dermal blood content. Figure 6C shows total percentage change between smallest and largest dermal thickness is 41.4%. In Figure 6D, percentage change between shallowest and deepest arterial depth is 32.1%. Similar trends of signal degradation are seen across the four different changes which occur with obesity: trans-epidermal water loss, blood perfusion, dermal thickness, and radial artery depth. Each scenario shows changes isolated to each feature of obesity. Figure 7 highlights changes seen at four different skin tones classified by epidermal melanin concentration. Total AC to DC signal ratio change is seen at 17.1% between 3% epidermal melanin concentration and 42% epidermal melanin concentration.

**Figure 6.**
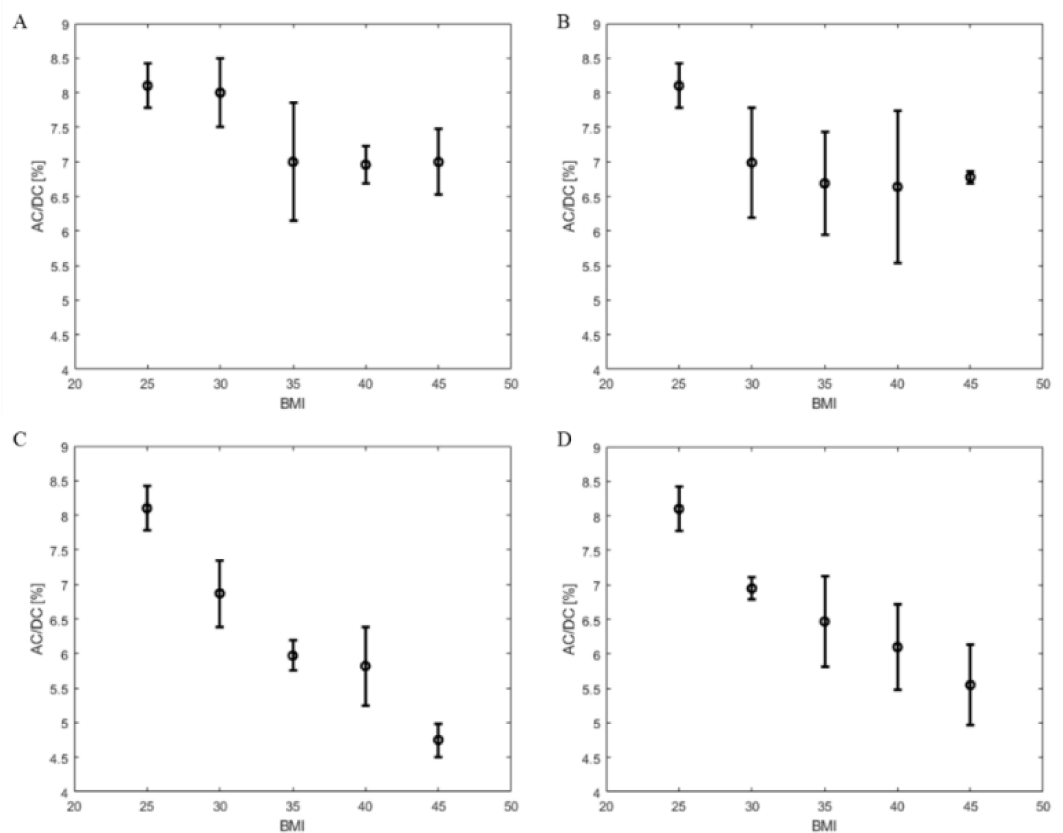
AD/DC PPG signal ratio for changes to trans-epidermal water loss (A), dermal blood content (B), dermal thickness (C), and radial artery depth (D) with BMI.

**Figure 7.**
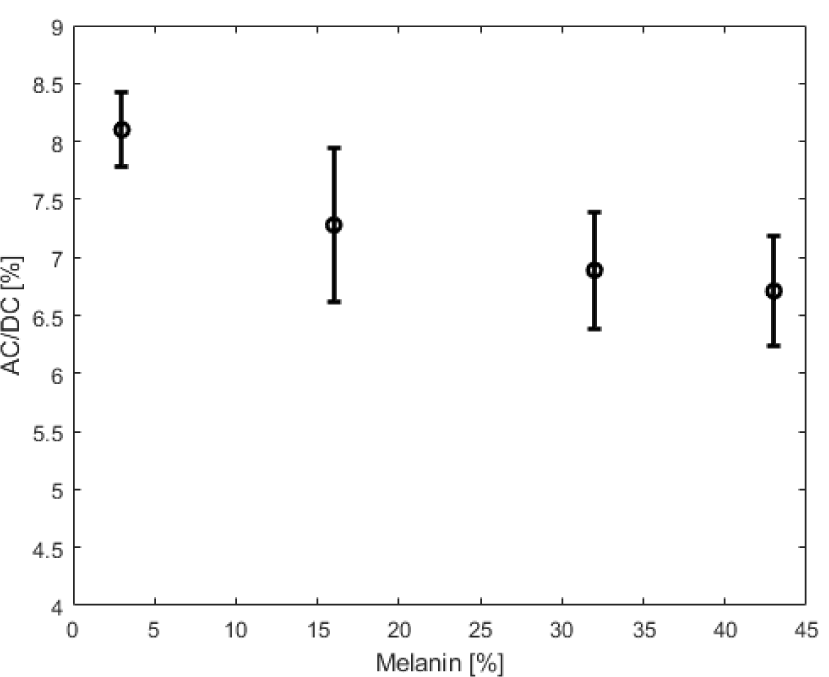
PPG AC/DC ratio change with melanin concentration change in epidermal layer for non-obese case.

Waveforms for different levels of obesity are generated through varying the combined dermal thickness and arterial depth to indicate corresponding physiological changes. Figure 8 shows generated waveforms at non-obese, obese 1, obese 2 and obese 3 at two different photon counts. Changes to BMI and obesity level is presented through the increase of dermal thickness from 1.0mm to 2.5mm and arterial depth from 2.5mm to 4.5mm. A comparison of the waveform shows the degradation of the baseline signal as well as changes to the wave shape itself. Across each level of obesity, at both 100 million and one billion photon counts, there is a significant diminishing of the amplitude of the signal. Figure 9A shows the four different signals superimposed on top of each other on the same curve. Baseline signal change between each obesity level and non-obese is shown in Table 2. The baseline change ranges from 12.7% to 54.5%. Similar changes are seen in both the 100 million photons set as well as the one billion photons set as to be expected. The advantage of using lower number of photons is the computational time for each simulation in exchange for a higher standard deviation at each point.

**Figure 8.**
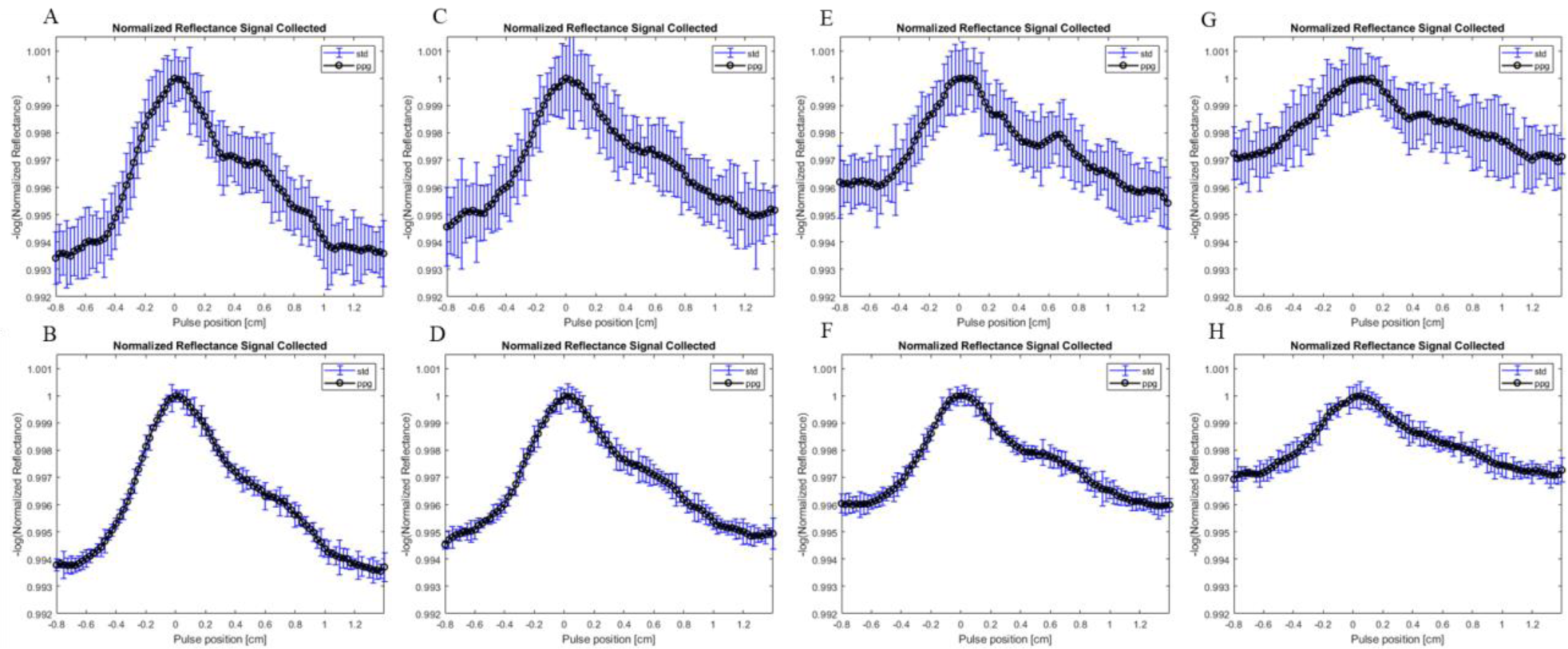
Variously generated waveforms at 660nm for different photon counts and different geometry representative of BMI change. (A) Non-obese waveform generated at 100 million photons and 15 trials. (B) Non-obese waveform generated at 1 billion photons and 5 trials. (C) Obese 1 waveform at 100 million photons and (D) 1 billion photons. (E) Obese 2 waveform at 100 million photons and (F) 1 billion photons. (H) Obese 3 waveform at 100 million photons and (G) 1 billion photons.

**Figure 9.**
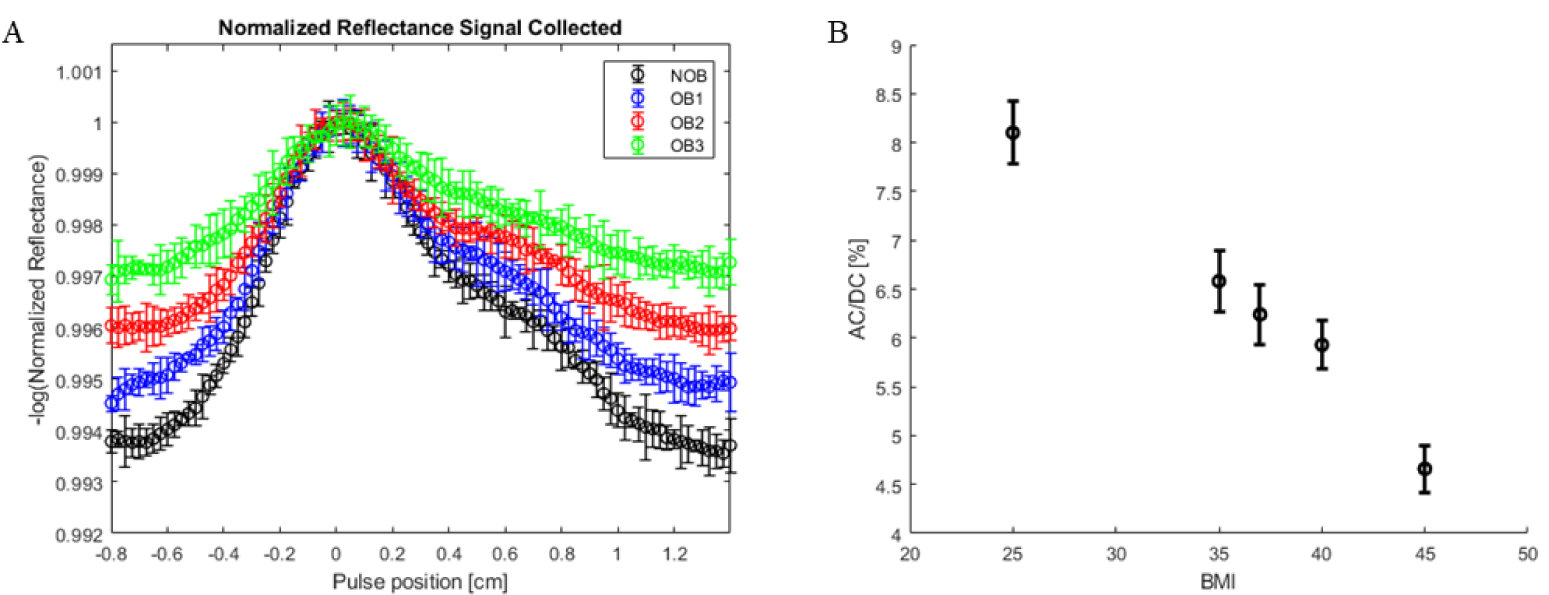
(A) Superposition of four different waveforms at 1 billion photons. Black: Non-Obese. Blue: Obese 1. Red: Obese 2. Green: Obese 3. (B) AC/DC ratio for each waveform signal comparison.

The non-obese BMI of 25kg/m^2^, the AC/DC signal ratio is 8.1% and at the maximum level calculated the ratio is 4.66%. This accumulates to the total percentage change of 43% between the Non-Obese waveform and Obese 3 waveform.

The degradation of the return signal gives an indication to the decrease in quality of the signal that will be received by the PPG device. A loss in signal quality is shown to increase as BMI and obesity level increase with changes in dermal thickness and arterial depth. PPG devices designed to periodically or continuously probe arterial blood pressure as a wearable device at the wrist must be designed to be mindful of the significant effects of obesity on optical performance. The advantage of the development of synthetic waveform is understanding these changes and the significance of their impact.

## 4. Discussion

We have demonstrated an approach to modeling PPG signals utilizing a combination of FEM and Monte Carlo modeling. Ultimately, our intent is to develop a wearable health device capable of assessing continuous cuff-less blood pressure with comparable level of performance across populations of varying body size. Cuff-less, continuous blood pressure monitoring can provide critical information concerning nocturnal cardiovascular function, which is a significant indicator of overall cardiac health. This work focuses on a commercial PPG sensor measurement of pulsatile flow in the radial artery.

Our simulations demonstrate that with no changes to the underlying vessel mechanical behavior, an increase in BMI significantly impacts the PPG waveform. The AC/DC ratio changes from 14% to 41% as BMI increases from 20 to 40 for isolated changes attributed with increasing level of obesity and BMI. Increase to dermal thickness increases the total volume of absorbers and scatterers through which photons must traverse to reach the artery and return to the detector. Increased radial artery depth also increases the total travel distance for photons to reach the vessel wall as and return to detector.

Vessel expansion and contraction, under pressure changes of the cardiac cycle, is directly associated with the pulsatility of the PPG waveform^59^, the presence of obesity degrades the quality of the signal even when the vessel mechanics remain unchanged. Synthetic waveform generation provides an insight and prediction of optical device functionality.

Future work will focus on the understanding of waveform morphology through connecting various cardiovascular conditions with the corresponding PPG. The addition of synthetic waveform generation can allow for study of isolated effects from a variety of factors on the output PPG. Post-processing of the PPG such as first and second derivative and frequency domain information can be used as tools for feature extraction for desired information.

This work is limited to the detection of the radial artery and does not consider the pulsatile effect of the superficial arterioles, this will be the focus of future addition to the present model.

